# Napping alters Functional Brain Responses in the Aged

**DOI:** 10.1101/2025.06.10.657597

**Authors:** Mathilde Reyt, Marine Dourte, Stella DeHaan, Michele Deantoni, Marion Baillet, Alexia Lesoinne, Sophie Laloux, Gregory Hammad, Fabienne Collette, Philippe Peigneux, Vincenzo Muto, Gilles Vandewalle, Mohamed Ali Bahri, Christophe Phillips, Christina Schmidt

## Abstract

Circadian rhythms shape the temporal organization of sleep and wakefulness and evolve throughout the adult lifespan, leading to higher sleep-wake cycle fragmentation with ageing. The increasing prevalence of daytime napping represents a visible manifestation of such fragmentation and has been suggested to forecast age-related cognitive decline. Here, we assessed the impact of napping on functional brain correlates of performance on a Sternberg working memory (WM) task using functional magnetic resonance imaging in 60 healthy older individuals, prospectively recruited with respect to their napping habits (39 females, age: 59–82y). As compared to non-nappers, nappers showed reduced hemispheric asymmetry in the dorsolateral prefrontal cortex (DLPFC, p< 0.001) and decreased performance at high WM load levels. Only in non-nappers was increased ipsilateral activation in the DLPFC associated with better performance at high WM load levels (p < 0.05), while contralateral activation across all WM load levels was not associated with better performance. These findings indicate that functional brain compensation and dedifferentiation processes vary according to an individual’s napping phenotype, potentially serving as a marker of inter-individual differences in cognitive and brain aging.

## Introduction

Large-scale population-based studies revealed that inter-individual variability characterizes the aging process as well as its associated cognitive decline (Lindenberger, 2014; Lupien & Wan, 2004; Nyberg et al., 2012). Increased group variance in cognitive processes implies that some individuals show very poor performance (up to pathological aging), whereas others maintain high levels of performance (Lupien & Wan, 2004). Poor performance has attracted much interest as it might be indicative of forthcoming disease, such as Alzheimer’s dementia. However, staying cognitively sharp at older age is not an exception. Successful agers are characterized by performance levels comparable to younger adults and keeping superior performance levels over time within their age-group (Lupien & Wan, 2004; Nyberg et al., 2012). Evidence from structural and functional neuroimaging are increasingly integrated to explain how the combined effects of adverse and compensatory neural processes produce both variable levels of cognitive change over the aging trajectory (Cabeza et al., 2018; Stern et al., 2020). Within this context, it has been suggested that high performing older adults recruit extra neural supply to allow maintenance of cognitive performance, despite an apparent age-related reduction in available neural resources, a concept referred to as compensation (Festini et al., 2018). Working memory (WM)-related brain activity has been particularly studied within this context. For instance, it has been observed that young adults show unilateral activation at low WM load but bilateral activation at high WM load in fronto-parietal networks, indicating the recruitment of supplemental brain areas to meet task demands (Bennett et al., 2013; Cappell et al., 2010; Mattay et al., 2006; Nyberg et al., 2009; Reuter-Lorenz et al., 2000; Schneider-Garces et al., 2010). On the other hand, older adults appear to already recruit bilateral neural resources in the dorsolateral prefrontal cortex (DLPFC) already at lower cognitive load levels, and would therefore have fewer supplemental or compensatory resources available at higher WM loads in order to adapt to the increasing task demands (Bennett et al., 2013; Cappell et al., 2010; Mattay et al., 2006; Nyberg et al., 2009; Reuter-Lorenz et al., 2000; Schneider-Garces et al., 2010). Besides the supplemental recruitment of contralateral frontal areas, a shift between relative activity levels in anterior and posterior brain areas is also observed (Davis et al., 2008), such that older adults show less activation of posterior brain regions and higher activation in anterior brain regions, compared to young adults. Importantly, literature reviews suggest that the manner and success of compensation is modulated by lifestyle variables, referred to as proxies of cognitive reserve (Cabeza et al., 2018; Stern et al., 2020). Life-course factors, such as stress, physical activity or intellectual engagement have been shown to contribute to the developmental progress of brain structure and function, as well as cognition over time (Dixon & Lachman, 2019; Reuter-Lorenz & Park, 2014).

How a fundamental mechanism such as sleep regulation impacts on age-related changes in cognitive processes remains largely underexplored. Aging goes along with changes in sleep timing, duration and structure (Taillard, Gronfier, et al., 2021), as well as an overall increased sleep-wake cycle fragmentation (Lim et al., 2011; Luik et al., 2013; Zuurbier et al., 2015). The latter has been associated with worsened cognitive performance (Luik et al., 2015; Oosterman et al., 2009), including WM (Lim et al., 2012). In the same vein, a particular emphasis has been attributed to the impact of habitual napping on cognitive performance in aging. Napping habit, reported in questionnaire or objectively quantified, using actimetry has been linked to decreased cognitive performance (Blackwell et al., 2006; Leng et al., 2019; Owusu et al., 2019; Reyt et al., 2022) and suggested to constitute a health risk factor for the prognosis and progression of Alzheimer’s disease (Li et al. 2022). Habitual napping in the aged is far from being uncommon and was reported to increase with advancing age such that more than half of adults aged 70 and above nap at least twice a week (Leng et al. 2018; 2019; Li et al. 2017). Here, we argue that the increased incidence of napping in older people is at least in part attributed to altered circadian sleep regulation. The latter is timed to achieve consolidated periods of sleep during night-time and a continuous period of wakefulness during daytime (Dijk and Archer, 2009). Circadian rhythms thereby shape the temporal organisation of sleep and wakefulness leading to consolidated bout of sleep during night-time and a continuous period of wakefulness during the day. This temporal sleep-wakefulness organization evolves throughout the adult lifespan, leading to higher sleep-wake cycle fragmentation with ageing. The increasing occurrence of daytime napping becomes in turns the most visible manifestation of this fragmentation. Hence, napping may reflect an “index of circadian disruption” (Deantoni, Reyt et al., 2024) with putative deleterious effects on cognitive and brain function. Accordingly, we reported that frequent daytime rest in people older than 60 years was not only associated with an altered 24-h organization of the rest-activity cycle, but also with reduced episodic memory performance (Reyt et al., 2022).

Based on the fact that the temporal organization of sleep-wake states is fundamental to brain function and prone to disruption during the aging process, we therefore explored whether habitual daytime napping interferes with functional brain compensation, underlying inter-individual variability in cognitive performance. After in-depth cognitive, sleep and circadian phenotyping, 60 healthy and cognitively unimpaired older adults (59 – 82 years old), prospectively recruited with respect to their napping habits (30 self-reported regular nappers vs 30 matched self-reported non-nappers), performed a Sternberg WM task while undergoing functional Magnetic Resonance Imaging (fMRI). We hypothesised that, compared to non-nappers, habitual nappers would be not only characterized by worse WM performance, but also by more “old age-like” functional dynamics in the brain responses corresponding to enhanced recruitment, mainly of the DLPFC, at lower WM loads. Given its potential position at the interface between vigilance and cognition (Portas, 1998), we further explored whether groups differ in task-related thalamic activation.

## Results

### Participants

After an extensive eligibility check (see methods), 30 habitual nappers and 30 non-nappers were prospectively selected and individually matched with respect to age, gender, educational level and season of assessment. Participants were classified as habitual nappers when subjectively reporting to nap at least three times a week for at least 45 minutes. As a consequence, habitual nappers did report a higher napping frequency (X^2^ = 37.03, p< 0.001) and duration (X^2^ = 28.48, p< 0.001) in their sleep agendas. In addition, actigraphy-derived outcomes (Figure 1) further confirmed a significantly higher daytime rest frequency (W = 84.5, p< 0.001) and duration (t = −3.59, p< 0.001) in habitual nappers as compared to non-nappers. Groups did not significantly differ, however, with respect to age, self-reported gender, and many potential confounding factors (education, subjective sleep quality, chronotype, affective state) (Table 1, all p_s_> 0.05). Importantly, groups also did not significantly differ with respect to their global cognitive status (MMSE and PACC5 score, all p_s_>0.05). Despite exclusion of participants with indices ≤ 18 and ≥30kg/m^2^ at study entrance, regular nappers presented a higher body mass index (BMI), compared to non-nappers (t = −2.06, p< 0.05). Also, habitual nappers reported a higher habitual daytime sleepiness than non-nappers (Table 1, Epworth Sleepiness Scale (ESS), Johns, 1991: W = 264.5, p< 0.01), while they did not differ in their subjective state of sleepiness (p> 0.05, Table 1, Karolinska Sleepiness Scale (KSS); Akerstedt & Gillberg, 1990) assessed before performance on the WM task in the MRI scanner.

**Figure 1.**
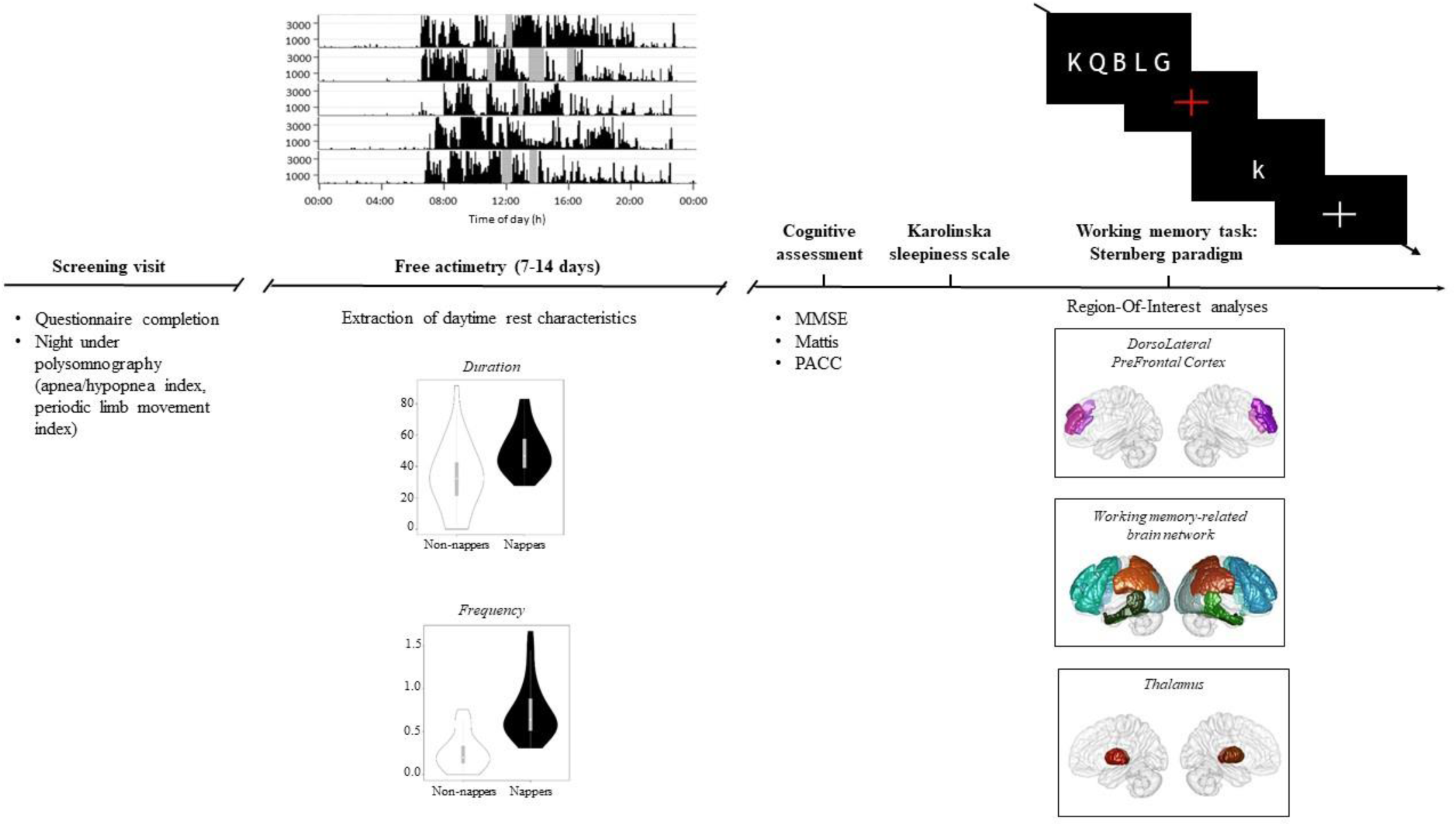
Schematic representation of the study protocol. After the screening visit, participants wore an actimeter during 7 to 14 days in order to evaluate daytime rest characteristics. Participant performed the Mini Mental State Examination (MMSE, Folstein et al, 1975), the Mattis Dementia Rating scale (Mattis, 1976) and the Preclinical Alzheimer’s Composite Score (PACC, Papp et al, 2017). After completing a Karolinska sleepiness scale (Akerstedt & Gillberg, 1990b), participants performed a Sternberg task in a 3T MRI scanner. The BOLD signal was then extracted from the dorsolateral prefrontal cortex, an extended working memory related brain network (sub-divided in frontal (blue), temporal (green), parietal (orange) and occipital (grey) regions) and the thalamus.

**Table 1.**
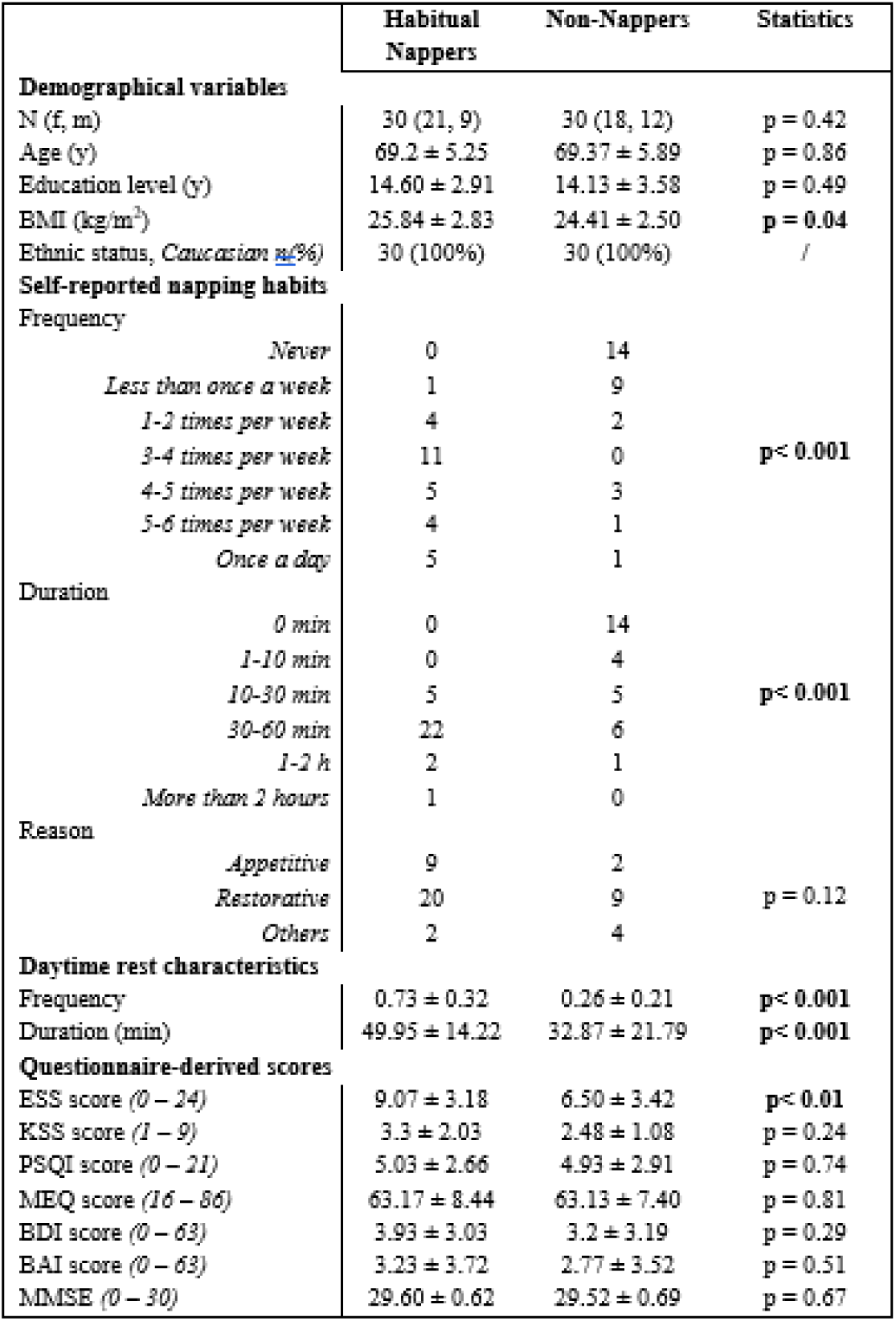
Demographical, self-reported napping habits, actigraphy-derived and questionnaire data by group (mean ± standard deviation). Note: f = female; m = male; y = years; BMI = Body Mass Index; ESS = Epworth Sleepiness Scale (Johns, 1991b); KSS = Karolinska Sleepiness Scale (Akerstedt & Gillberg, 1990); PSQI = Pittsburgh Sleep Quality Index (D J Buysse et al., 1989a); MEQ = Morningness-Eveningness Questionnaire (Horne and Östberg, 1976); BDI = Beck Depression Inventory (Beck et al., 1961); BAI = Beck Anxiety Inventory (Beck et al., 1988); MMSE = Mini-Mental State Examination (Folstein et al, 1975)

### Napping affects WM performance at high load levels

Working memory performance was assessed with the Sternberg WM paradigm (Figure 1; Sternberg, 1966), that allows for a parametric modulation of working memory load and has been extensively studied using fMRI. Participants had to indicate whether a probe letter was part of a series of previously presented letters. WM load or Memory Set Size (MSS) corresponds to the number of letters presented during the encoding phase (2-7 letters, see methods). We first assessed the impact of WM load, nap group as well as their interaction on task performance using generalised linear mixed models (GLMM), including BMI and daytime sleepiness as covariates. According to the Akaike criterion (see methods for model comparisons), the model assessing performance accuracy also included demographical variables (age, sex and educational level).

As expected, higher WM load was associated with decreasing accuracy from MSS 5 onwards (all p<0.01, Table 2, Figure 2) and increased reaction times from MSS 2 onwards (all p< 0.001, Table 2, Figure 2). Significant interactions for both d’ and reaction times revealed that, as compared to habitual nappers, non-nappers performed faster (t= 2.31, p< 0.05, Figure 2) and better (t= −1.98, p=0.048, Figure 2) at MSS 7.

**Figure 2.**
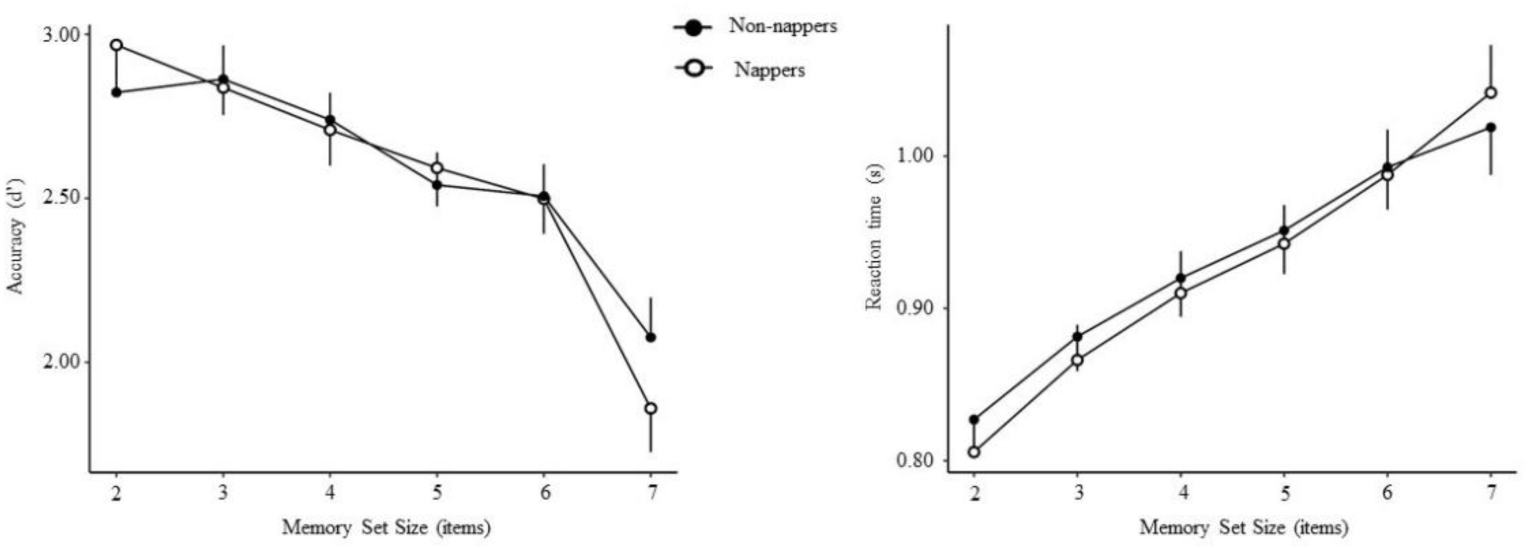
Mean accuracy (d’ measure, on the left) and median reaction times (on the right) across memory set sizes for non-nappers (black circles) and habitual nappers (white circles) (mean ± SEM).

**Table 2.**
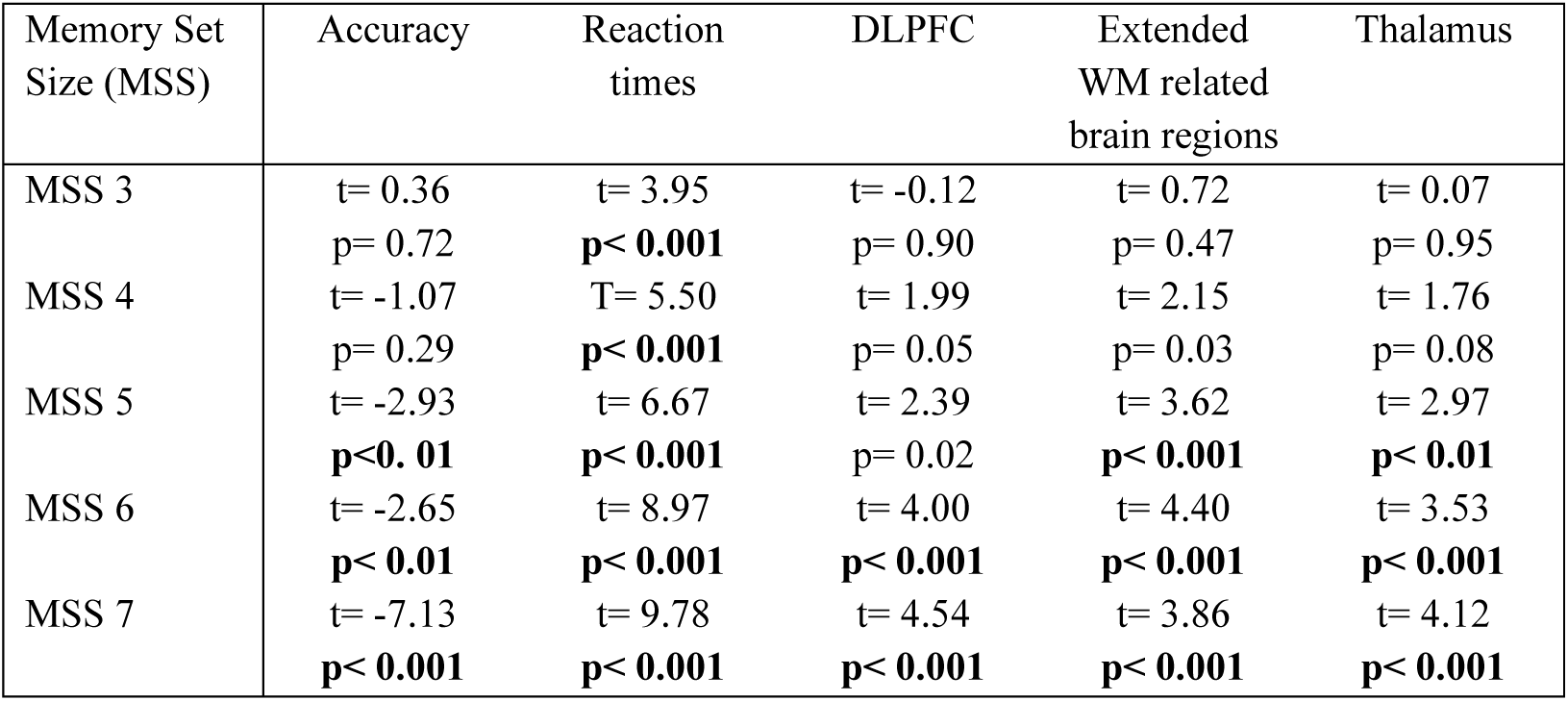
Effect of working memory load during the Sternberg task on accuracy, reaction times and BOLD signal in the three regions of interest (Helmert coding). Note: WM = working memory

In addition, accuracy decreased with increasing age (t= −2.87, p< 0.01) and women showed lower performance compared to men (p = 0.04).

### Habitual napping affects the functional brain correlates underlying WM

Brain activitation was operationalized using the BOLD signal derived from functional MRI during the performance of the Sternberg paradigm. The involvement of the dorsolateral prefrontal cortex (DLPFC), particularly Brodmann’s area (BA) 9/46 in this WM task has been crucially reported (Haque et al., 2021) to be dynamically recruited over increasing WM load (Altamura et al., 2007; Bennett et al., 2013; Cappell et al., 2010; Höller-Wallscheid et al., 2017; Schneider-Garces et al., 2010).

Generalised linear mixed models were first computed to assess whether BOLD activation in the DLPFC is modulated by WM load (2-7 letters, Helmert contrasts), group (habitual nappers vs non-nappers, treatment contrast: reference = non-nappers), hemisphere (left vs right, treatment contrast: reference = left), as well as their interactions. Laterality was included as a factor, considering that compared to younger adults, older adults have been reported to show more bilateral activation (Bennett et al., 2013; Cappell et al., 2010; Mattay et al., 2006; Nyberg et al., 2009; Reuter-Lorenz et al., 2000; Schneider-Garces et al., 2010) and putatively less differentiated hemispheric recruitment (Cabeza, 2002). Finally, BMI, daytime sleepiness and regional volume were included as covariates. Higher WM load was associated with a higher BOLD signal in the DLPFC (BA 9/46) starting at MSS 6 (p< 0.001; Table 2, Figure 3). Furthermore, a significant laterality*group interaction revealed that, independently of WM load and as compared to non-nappers, regular nappers presented higher BOLD activation in the right DLPFC (t= 3.35, p< 0.001, Figure 3), while no significant differences were observed in the left DLPFC.

**Figure 3.**
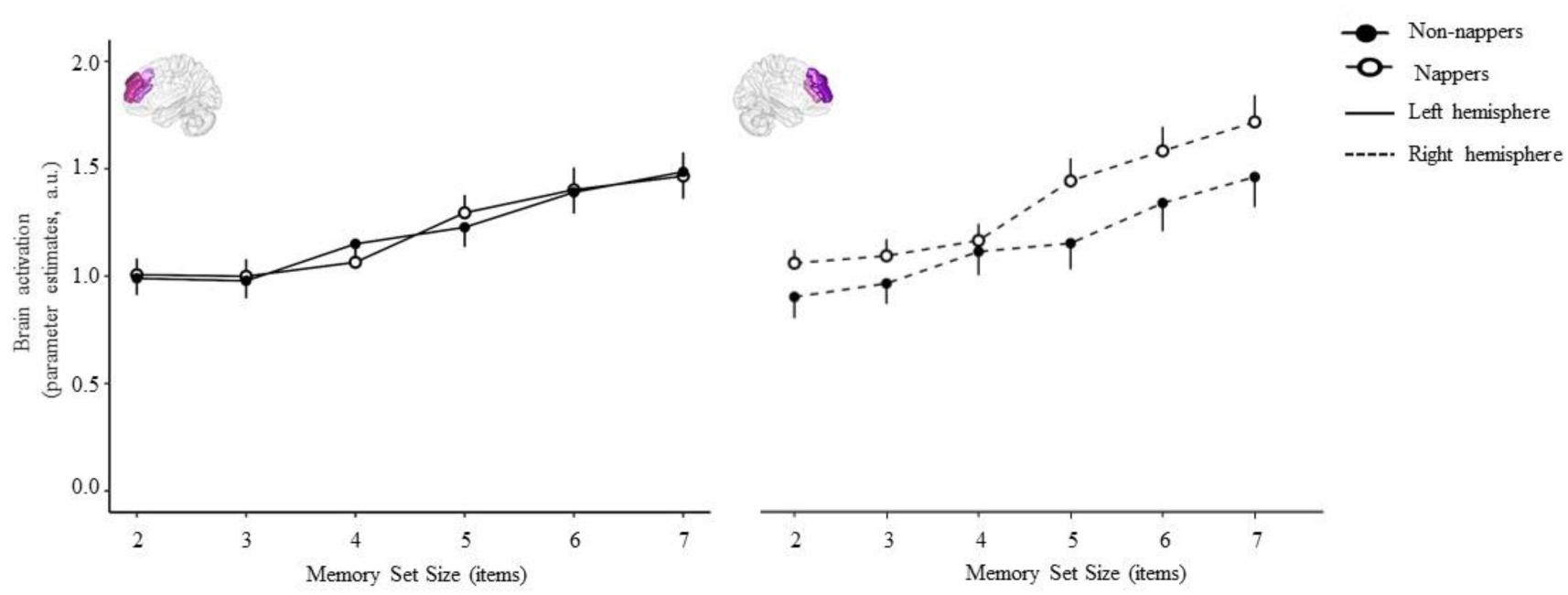
Mean BOLD signal across memory set sizes for the left dorsolateral prefrontal cortex (DLPFC) (on the left) and right DLPFC (on the right, dotted lines) for non-nappers (black circles) and habitual nappers (white circles) (mean ± SEM).

Some studies highlighted that hemispheric asymmetry reduction observed in older adults could not be explained by compensation-based models (McDonough et al., 2013; Roe et al., 2020) but may rather reflect less specific neural responses, also-called dedifferentiation (Knights et al., 2021; Morcom & Henson, 2018). Age-related bilateral activation during a Sternberg task is indeed consistent with both dedifferentiation and compensation-based models (Carp et al., 2010). One requirement to define compensation is that increased BOLD activation is linked to performance improvement (Cabeza et al., 2018). If hemispheric asymmetry reduction in nappers is compensatory, WM performance should therefore be explained by BOLD signal in the DLPFC. We thus performed GLMM to test whether task performance (d’ or reaction time) was associated with WM load (2-7 letters, Helmert contrasts), group (habitual nappers vs non-nappers, treatment contrast: reference = non-nappers), BOLD signal as well as their interactions. Daytime sleepiness and BMI were added as covariates and, according to Akaike criterion (see methods for model comparisons), model assessing accuracy also included age, gender and educational level. An WM load*BOLD signal*group interaction effect revealed that higher BOLD signal in the left DLPFC at MSS 7 was associated with accuracy only in non-nappers (t= −2.00, p= 0.047). No significant results were observed for the right hemisphere and reaction times (all p_s_> 0.05).

### Habitual napping & regional specificity in BOLD signal dynamics underlying working memory performance?

Studies investigating age differences in compensation-related processes have reported recruitment of supplemental brain areas in a more widespread WM-related brain network (Cappell et al., 2010; Schneider-Garces et al., 2010). Accordingly, region of interest (ROI) analyses were conducted on task-related BOLD signals in the DLPFC, but also in the frontal, parietal, temporal and occipital regions (extended WM-related brain network ROI, see methods, Figure 1). Besides differences in bilateral recruitment, compensatory-based models also posit a posterior-anterior shift in brain activation during WM performance, such that increased activation in anterior brain regions and reduced activity in posterior brain regions are associated with better performance in older adults (e.g. Davis et al., 2008; Festini et al., 2018). To assess whether napping habits affects brain activity in this framework, we extracted the BOLD signal in the ROIs encompassing the frontal, parietal, temporal and occipital lobes in the WM-network previously reported to be recruited by older adults during the Sternberg task (Cappell et al., 2010; Schneider-Garces et al., 2010, see also methods). GLMMs were computed to assess whether brain activity is modulated by WM load (2-7 letters, Helmert contrasts), group (nappers vs non-nappers, treatment contrast: reference = non-nappers), regions (frontal, parietal, temporal, occipital, treatment contrast: reference: frontal) and hemisphere (left vs right, treatment contrast: reference = left), with covariates BMI, daytime sleepiness and regional volume. Increasing WM was associated with higher BOLD signal starting at MSS 5 (all p<0.001, Table 2, Figure 4). Overall, the left hemisphere was more activated (t= −13.79, p<0.001) than the right hemisphere (Figure 4). Compared to the frontal region, BOLD signal was higher in parietal (t= 5.79, p< 0.001, Figure 4), temporal (t= 4.61, p< 0.001, Figure 4) and occipital regions (t= 11.87, p< 0.001, Figure 4). Hemisphere*regions interaction effects revealed higher activation in the left parietal (t= 5.27, p< 0.001, Figure 4), left temporal (t=6.40, p< 0.001, Figure 4) and left occipital lobe (t= 7.86, p< 0.001, Figure 4). Furthermore, as compared to non-nappers, BOLD signal was higher in habitual nappers in parietal than in frontal regions (interaction group*regions: t= −1.97, p = 0.048, Figure 4).

**Figure 4.**
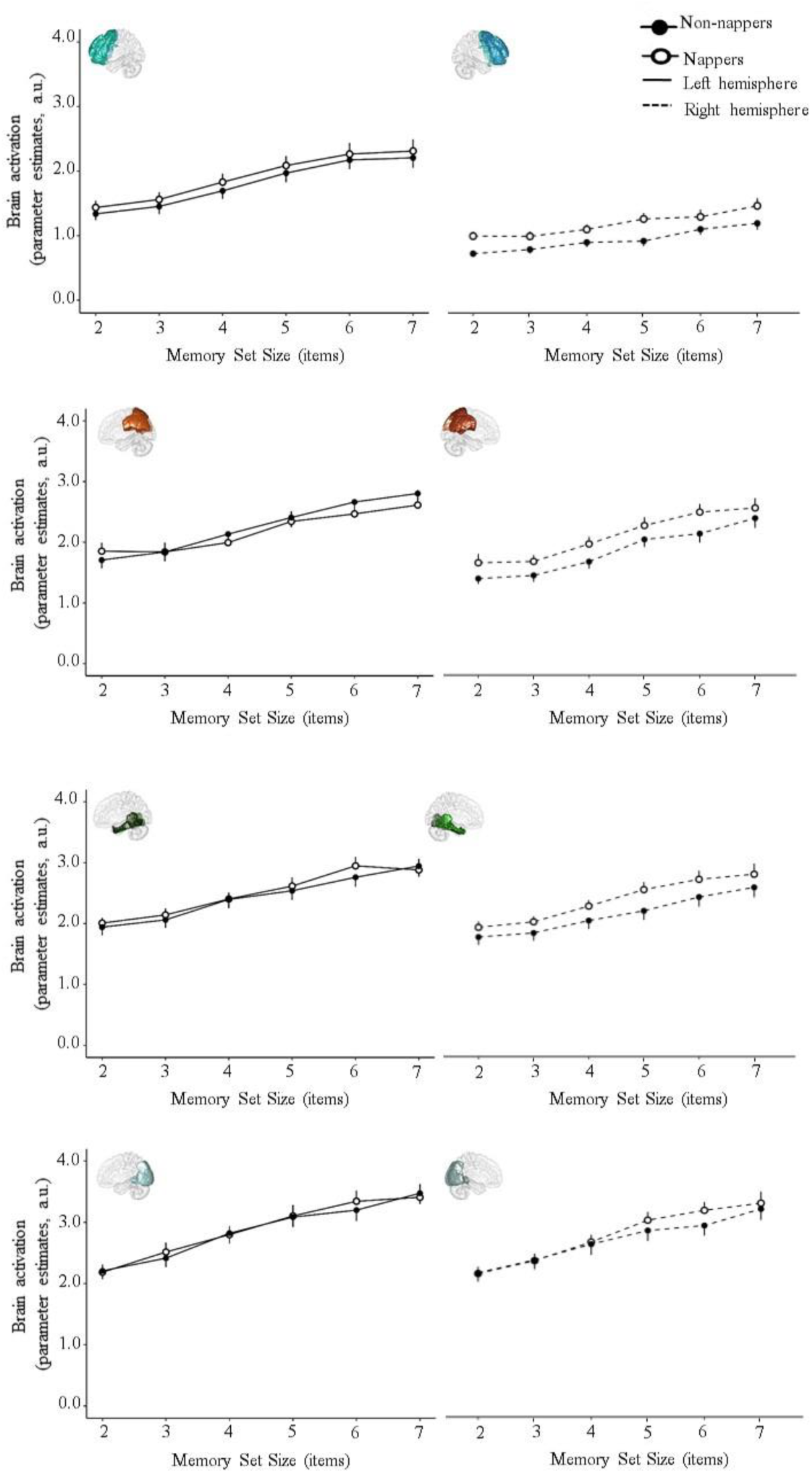
Mean BOLD signal across memory set sizes for the frontal, parietal, temporal and occipital regions for non-nappers (black circles) and habitual nappers (white circles) (mean ± SEM). Continuous lines correspond to left hemisphere and dotted lines represent the right hemisphere.

### Habitual napping & hemispheric asymmetry reduction

Finally, a separate ROI analysis was performed by extracting BOLD signal in the thalamus (Figure 1). Task-evoked thalamic BOLD activation has been shown to play a mediating function in the interaction between attention and arousal (Portas, 1998). Likewise, responses have been suggested to counter arousal-modulating effects in response to sleep loss or circadian misalignment (e.g. Muto et al. 2016), as may be expected in nappers. Building on our previous findings, showing altered circadian sleep regulation and vigilance modulation in habitual nappers (Deantoni et al., 2023), we performed GLMM to assess whether thalamic BOLD activation is modulated by WM load (2-7 letters, Helmert contrasts), group (habitual nappers vs non-nappers, treatment contrast: reference = no-nappers), and hemisphere (left vs right, treatment contrast: reference = left). As expected, a significant effect of WM load was observed in the thalamus with higher BOLD activation starting at MSS 5 (all p<0.01, Table 2, Figure 5). Interestingly, as compared to non-nappers, habitual nappers activated more the thalamus over all WM and both hemispheres (t= 2.17, p = 0.03, Figure 5). Finally, while the right thalamus was overall more activated, compared to the left thalamus (t = 14.33, p< 0.001, Figure 5), a significant group*hemisphere interaction revealed that the BOLD signal in the right hemisphere was higher in regular nappers, compared to non-nappers (t = 3.53, p< 0.001, Figure 5).

**Figure 5.**
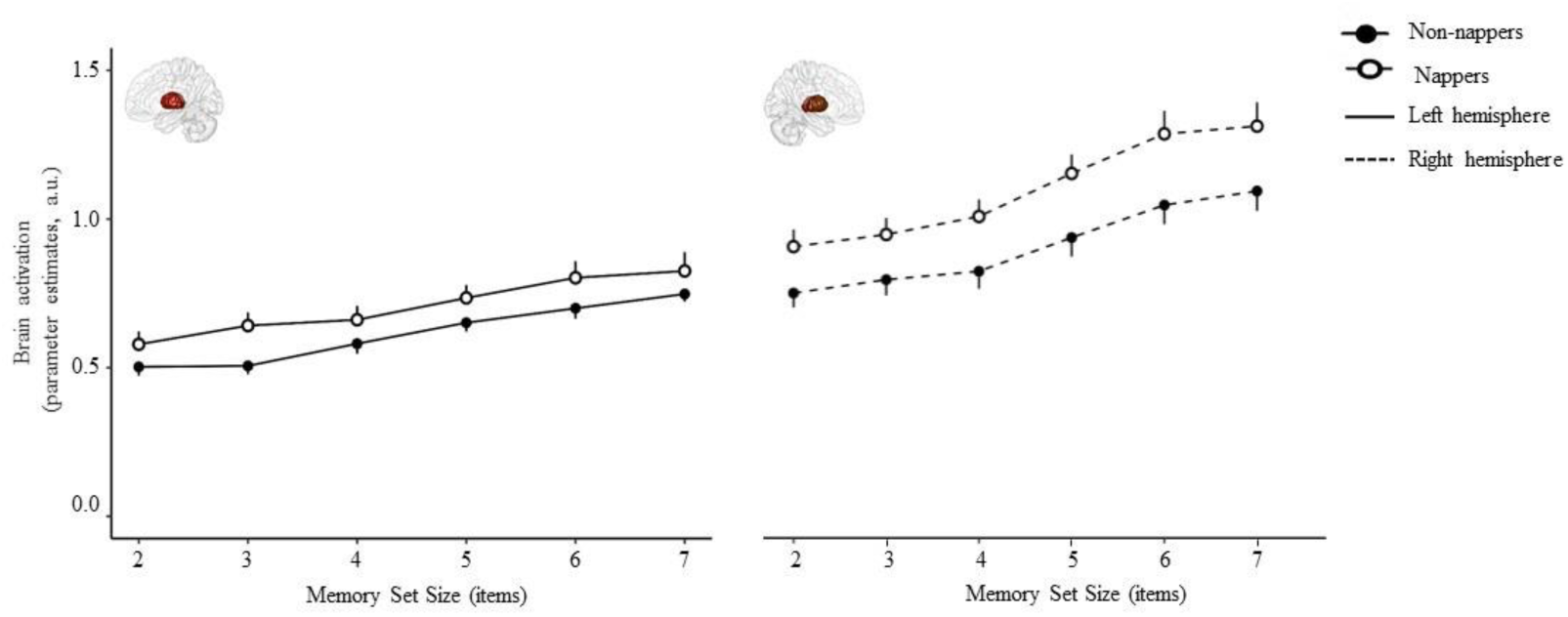
Mean BOLD signal across memory set sizes for the left thalamus (on the left) and right thalamus (on the right, dotted lines) for non-nappers (black circles) and habitual nappers (white circles) (mean ± SEM).

## Discussion

The older proportion of the population is increasing worldwide (Lindenberger, 2014), and cognitive decline represents a prominent factor for determining the health span of an older and aging population. In the absence of a strong genetic program (Morris et al., 2019), person-specific endowments, environmental opportunities and constraints act as modulators and modifiers on cognitive and brain aging (Cabeza et al., 2018; Reuter-Lorenz & Park, 2014; Stern et al., 2020). The sleep-wake cycle and its age-related fragmentation reflect potentially malleable life-style components for cognitive aging (Lim et al., 2012; Luik et al., 2015; Oosterman et al., 2009). The increased incidence of napping and/or daytime rest has been recently suggested to be associated with or even forecast cognitive decline in longitudinal settings (Leng, 2019, Li, 2022, Reyt et al. 2022). This aligns with growing epidemiological evidence linking habitual napping to health risk factor in older adults (Li et al., 2022; Sun et al., 2022). Additionnaly, we highlighted altered circadian regulation in regular nappers (Deantoni, Reyt, et al., 2024), suggesting that such disruptions may contribute to these negative cognitive and health-related outcomes. Besides behavioral differences detectable at high working memory load, our results provide first evidence of altered WM-related brain responses in healthy older habitual nappers, most strongly expressed as a specifically reduced hemispheric asymmetry in the DLPFC.

The Compensation-Related Utilization of Neural Circuits Hypothesis (CRUNCH; Reuter-Lorenz and Cappell, 2008) has been developed based on age-related changes in behavioural and cerebral responses to increased WM load. This model predicts higher brain activation with increasing WM load to meet task demands. In line with this prediction, we observed increased BOLD activation with increasing WM load in all ROIs and in both groups. Working memory processes critically involve the DLPFC, and more particularly the left DLPFC (Haque et al., 2021). Furthermore, the CRUNCH model posits that age-related brain alterations result in an over-activation that can be expressed as bilateral activation (Cabeza, 2002). During the Sternberg task, at low cognitive loads, young adults showed an asymmetric recruitment in the ipsilateral (left) hemisphere whereas a bilateral activation was observed in older adults (Cappell et al., 2010; Schneider-Garces et al., 2010). Within this context, increased recruitment of the contralateral hemisphere in the DLPFC in habitual nappers suggests that they may be characterized by a more “old age-like” brain function. At the behavioural level, we observed that, as compared to non-nappers, regular nappers did not significantly differ in their performance (accuracy and reaction times), except for the highest WM load (MSS 7). Worse performance of nappers at the highest WM load indicates that they may have reached earlier their maximal WM capacity limit. This hypothesis is supported by a trend towards higher BOLD activation in the ipsilateral hemisphere in non-nappers at the highest WM load and higher BOLD signal in the contralateral hemisphere in habitual nappers, as compared to non-nappers. Whereas non-nappers might compensate with higher brain activity and thus maintain performance levels, performance decline in habitual nappers could be explained by reaching their maximal brain activity. Recruitment of additional resources associated with maintenance of performance are characteristics of compensatory process described in the context of successful aging. Our data might thus suggest that habitual nappers compensate at the functional level, until maximal WM capacity is reached and/or supplemental recruitment is no more possible, thereby resulting in performance decline (Cabeza & Dennis, 2012).

However, the analysis assessing the association between performance and BOLD signal does not indicate that the contralateral activation observed in regular nappers is linked to WM performance. Besides functional compensation, hemispheric asymmetry reduction in older adults has been assimilated to brain “dedifferentiation” (Knights et al., 2021; Morcom & Henson, 2018). This process refers to a reduced distinctiveness of neural responses, leading to less specific brain activation patterns. Single neuron-recordings in animals (Ding et al., 2017; Hua et al., 2006; Schmolesky et al., 2000) provide strong evidence for an age-dependent reduction in the neural selectivity, also defined as smaller differences between activity for the specific and non-specific stimuli of a brain region (Koen & Rugg, 2019). Notably, the nature of the task, its behavioural responses, as well as the BOLD signal dynamics in task-related brain regions should be considered to determine if hemispheric asymmetry reduction reflects compensation, dedifferentiation, or both. The absence of a significant association between performance and contralateral recruitment may suggest that hemispheric asymmetry reduction observed in the context of napping habits is not necessarily related to functional compensation. Nonetheless, an association between WM performance and higher BOLD signal at the highest WM load in the ipsilateral hemisphere might indicate functional compensation in non-nappers. Our study was not designed to differentiate between both processes, but compensation and dedifferentiation are not incompatible, and have been suggested to act simultaneously in a study investigating age-related changes in performance during a Sternberg paradigm (Carp et al., 2010). Actually, functional compensation is a process to counteract adverse effects of brain alterations, such as dedifferentiation.

The arousal-promoting thalamus is implicated in the maintenance of stable wake states and plays a key role in mediating the interaction between attention and arousal in humans (Portas, 1998, Sturm, 1999). Increased thalamic activity has also been observed with increasing cognitive demands (Schiff, 2008). Also, thalamic activity has been consistently shown to increase under conditions of sleep deprivation and circadian misalignment (Ma, 2015, Muto et al. 2016). Within this context, Maire et al. (2015), for example, observed that individuals more vulnerable to sleep deprivation need to recruit more the thalamus across the duration of the task, compared to behaviorally more resilient individuals (who exhibit less attentional lapses, reduced task disengagement). Here, we did not proceed to such a sleep manipulation, but observed that habitual nappers, characterized by more daytime rest bouts and increased overall sleepiness during daytime, also showed enhanced thalamic recruitment during task engagement. These results potentially point towards an increased wake state instability. In this respect, napping, as an intrusion of sleep into the active wake period, might be used as a strategy to counteract the inability of older adults to maintain wake states over extended periods of time (Carskadon et al., 1982). Additionally, the rising frequency and duration of napping in older adults may be partially attributed to altered circadian sleep regulation, as our recent findings reveal associations between napping habits and these disruptions. Habitual nappers also exhibited altered modulation in sleepiness and vigilance at specific circadian phases (Deantoni, Reyt, et al., 2024). Considering the thalamus’ role in regulating sleep-wake cycles, these circadian disruptions may impair thalamic activity, leading to sleep pattern alterations and potentially affecting cognitive performance. Whether these changes are specifically driven by circadian sleep and/or wake-promoting mechanisms remains to be assessed.

This is to the best of our knowledge the first study to explore the impact of habitual napping on the functional brain responses underlying a WM challenge. Our data indicate hemispheric asymmetry reduction in habitual nappers, compared to non-nappers, that may be underlined by processes of dedifferentiation and functional compensation, depending on behavioral performance. Notably, age-related structural brain changes are now considered in conceptual models of cognitive aging (Park & Reuter-Lorenz, 2009; Reuter-Lorenz & Park, 2014), positing that compensatory neural activation is an adaptive mechanism in response to decline, both in brain function and structure. Future studies should investigate the potential consequences of napping habits on brain structure and assess whether brain activity differences could also be underpinned by structural brain modifications, particularly in relation to cortical gyrification. Considering the association between circadian sleep regulation and cortical gyrification, as suggested by previous findings linking distinctive REM sleep allocation to regional brain structure changes in aging (Deantoni et al., 2023), it would be valuable to explore how these modifications may contribute to brain maintenance concepts derived from structural MRI (Cabeza et al., 2018; Stern et al., 2020). Dedifferentiation and functional compensation are dynamic and modifiable by lifestyle factors, suggesting that acting on protective factors, such as a good sleep hygiene, could enhance these mechanisms and thus improve cognitive performance. Our study thereby supports the need of longitudinal designs on napping habits during aging and to determine whether the restoration of a consolidated 24-h rest-activity pattern could lead to cognitive improvement and the generalization to a less selective and more representative sample of the general population.

## Methods

This cross-sectional study consisted of three phases: (1) a telephone interview and screening visit, (2) a pre-laboratory field actimetry and sleep diary study, and (3) a cognitive assessment followed by a functional MRI scanning during a WM task, collectively scheduled to the afternoon hours. Further details of the study procedure can be found in Reyt et al. (2022) and Deantoni, Reyt, et al. (2024).

### Participants

Recruitment aimed at covering a wide range of socio-economic classes and was performed via advertisement in newspapers, radio and university and by taking advantage of already existing GDPR-compliant databases at the research unit. All contacted volunteers were retired and lived independently at home. Seven hundred and seventy-three individuals aged ≥ 60 were initially contacted. Out of the sample, 166 passed the screening visit. None of the participants indicated moderate or severe depression (Beck Depression Inventory [BDI-II] < 19, Beck et al., 1961) or severe anxiety (Beck Anxiety Inventory [BAI] < 30, Beck et al., 1988). Clinical symptoms of cognitive impairment were assessed by the Mini Mental State Examination [MMSE score > 26] (Folstein et al., 1975) and the Mattis Dementia Rating scale [score > 130] (Mattis, 1976). Screening for major sleep disorders was performed during a night of polysomnography and 72 participants were excluded (apnea/hypopnea index and period limb movement index > 15/h) (for the final sample of 94 participants: mean apnea/hypopnea index: 5.48 ± 4.63, mean Periodic Limb Movement index: 3.07 ± 5.51). Other exclusion criteria included Body Mass Index (≤ 18 and ≥30 kg/m^2^), participants reporting a history of diagnosed psychiatric conditions or severe brain trauma, chronic medication affecting the central nervous system, diabetes, smoking, caffeine (> 4 cups/day), excessive alcohol (> 14 units/week) or other drug consumption, and travelling more than one time zone in the 3 months prior to the study begin. Participants with stable treatment (> 6 months) for hypertension and/or hypothyroidism were included in the study.

For this cross-sectional assessment, a group of 30 nappers and 30 non-nappers were selected out of the sample and matched at the group level with respect to age, gender, educational level and season of assessment. Demographic characteristics of the two groups are summarized Table 1. Participants reported their napping habits through a questionnaire. Daytime rest duration and frequency were retrospectively assessed through actimetry recordings in both groups (see also below). Habitual nappers were defined as individuals who napped at least three times per week for a minimum of 45 minutes, consistently for at least one year. The non-napping group included individuals who reported either not napping, napping only occasionally, or taking naps lasting for a maximum of 10 minutes.

The study was approved by the local Ethics Committee of the University Hospital and of the Faculty of Psychology, Speech Therapy and Educational Sciences at the University of Liège (Belgium) and performed in accordance with the Declaration of Helsinki. Participants gave written informed consent and received a financial compensation.

### Actigraphy: daytime rest characteristics

As described in Reyt et al. (2022), daytime rest habits were assessed over a period of at least 8 days (13.60±1.90 days, range 8-15) during which participants wore an actigraphic device (Motionwatch 8, CamNtech, UK) at the non-dominant wrist and completed a sleep diary. In this latter, participants were asked to report actigraph removal periods, which were visually identified and excluded from the analysis. Motor activity was aggregated into 30-sec activity counts and the Munich Actimetry Sleep Detection Algorithm (MASDA) (Roenneberg et al., 2015) was used to detect consolidated daytime rest periods. The performance of this algorithm against visual scoring was previously reported (Reyt et al., 2022). Actigraphy data were processed using the open-source software pyActigraphy (v1.0) (Hammad et al., 2021).

Daytime rest periods were identified during the biological day (see Reyt et al. (2022)) and their duration ranged from 15 minutes to 4 hours. Based on detected daytime rest periods, frequency was calculated as the mean number of these periods per day and duration defined as the overall mean duration.

### Questionnaires

Questionnaires were administered at study entrance and included the Pittsburgh Sleep Quality Index (PSQI, Buysse et al., 1989), the Epworth Sleepiness Scale (ESS, Johns, 1991), the Morningness-Eveningness Questionnaire (MEQ, Horne & Östberg, 1976), the Beck Depression Inventory (BDI-II, Beck et al., 1961) and the Beck Anxiety Inventory (BAI, Beck et al., 1988).

### Cognitive assessment

Before the fMRI scan, participants completed a cognitive assessment including the Mini-Mental State Examination (Folstein et al, 1975), the Mattis Dementia Rating scale (Mattis, 1976), as well as the Pre-clinical Alzheimer’s Disease Composite Score (PACC, Papp et al., 2017).

### Working memory paradigm

Data were usually acquired during the afternoon (between 13:30 and 16:00) to avoid confounding effects of time of day on cognitive performance and its underlying cerebral correlates.

Participants completed a Sternberg task including six memory set sizes (MSS) varying from 2 to 7 during a functional MRI acquisition session (see below). Stimuli were consonants (except letters P, W, X and Z due to their high distinctiveness). Letters were presented at the screen’s centre, in “Arial” font and a 120-pixel font size. Uppercase letters were displayed for encoding and lowercase letters were used for probes.

During encoding, a set of 2 to 7 white letters were simultaneously displayed on a black screen (Figure 1). Each trial started by the presentation of a memory set during 3 sec, followed by a 1 sec maintenance interval (red fixation cross). At retrieval, the probe letter was presented during 0.8 sec, followed by a white fixation cross during 1.2 sec. During the retrieval period (2 sec), participants had to indicate as quickly and accurately as possible, using one of the two responses keys, whether the probe was part of the memory set or not. Each block was composed of 4 trials of the same MSS and participants completed 48 blocks (8 blocks by 6 MSS). A grey fixation cross was displayed during 15 sec between each block. The order of block presentation was pseudo-randomized to avoid presentation of consecutive blocks of the same MSS. For each MSS, the probe was part of the memory set on half of the trials.

Before being installed in the MRI scanner, participants underwent a training session, composed of 10 trials using the 6 MSS. The training was performed as many times as necessary to ensure that participants understood the task and reached at least 70% accuracy. Five trials were also performed just after participants’ installation in the MRI scanner to verify response keys functionality.

The Sternberg task was presented using Psychophysics Toolbox Version 3.0.15 (Brainard, 1997; Kleiner et al., 2007; Pelli, 1997) running on Matlab R2017a (The MathWorks, Inc., Natick, MA).

### Behavioural data analysis

Generalised linear mixed models (GLMM) were conducted to assess the effect of group (habitual nappers vs non-nappers, reference = non-nappers), memory load (2-7 letters) as well as their interaction on WM accuracy (d’; Stanislaw & Todorov, 1999) and reaction times, separately. Group and memory load were defined as categorical fixed effects and subjects as random effect. Performance at each MSS was compared to the mean performance of all previous MSS (Helmert coding). Non-nappers were defined as the reference group for the treatment contrast. In order to determine whether demographical variables, such as age, gender and educational level, should be considered as covariates in GLMM, we compared models with and without these variables. The comparison was performed in R version 3.6.3 (R Core Team, 2020), requiring the package AICcmodavg (Mazerolle, 2020). The model with the lower Akaike criterion was selected. After model comparisons, age, sex and education had to be included as covariates for WM performance, but not for reaction times. Daytime sleepiness and BMI were also included as covariates since group significantly differ on these variables. Significance was based on a p-value< 0.05. Statistical data analyses for performance measures were conducted on R version 3.6.3 (R Core Team, 2020) using packages stats (R Core Team, 2020), tidyr (Wickham et al., 2019), lme4 (Bates et al., 2015), lmerTest (Kuznetsova et al., 2017) and Rmisc (https://www.rdocumentation.org/packages/Rmisc/versions/1.5.1).

### Functional magnetic resonance imaging protocol

Functional MRI time series were acquired on a whole-body 3T scanner (Magnetom Prisma, Siemens Medical Solutions, Erlangen, Germany) operated with a 64-channel receiver head coil. Multislice T2*-weighted functional images were acquired with the multi-band gradient-echo echo-planar imaging sequence (CMRR, University of Minnesota) using axial slice orientation and covering the whole brain (36 slices, multiband factor = 2, FoV = 216×216 mm², voxel size 3×3×3 mm³, 25% inter-slice gap, matrix size 72×72×36, TR = 1170 ms, TE = 30 ms, FA = 90°). The four initial volumes were discarded to avoid T1 saturation effects. During fMRI acquisition, cardiac pulse and respiration were recorded with a pulse oximeter and respiration belt. A gradient-recalled sequence was applied to acquire two complex images with different echo times (TE = 10.00 and 12.46 ms respectively) and generate field maps for distortion correction of the echo-planar images (EPI) (TR = 761 ms, FoV = 192×192 mm², 64×64 matrix, 48 transverse slices (3 mm thickness, 50% inter-slice gap), flip angle = 90°, bandwidth = 260 Hz/pixel). For anatomical reference, a high-resolution T1-weighted image was acquired for each subject (T1-weighted 3D magnetization-prepared rapid gradient echo (MPRAGE) sequence, TR = 1900 ms, TE = 2.19 ms, inversion time (TI) = 900 ms, FoV = 256×240 mm², matrix size = 256×240×224, voxel size = 1×1×1 mm³, acceleration factor in phase-encoding direction R=2).

### fMRI data: pre-processing and analysis

Image processing and analysis were performed using the SPM12 software (Centre for Human Neuroimaging, www.fil.ion.ucl.ac.uk/spm) running on Matlab R2017a. EPI time series were reoriented into the MNI space using the SPM template, then corrected for motion and distortion using the “Realign and Unwarp” module (Andersson et al., 2001) together with the FieldMap toolbox (Hutton et al., 2002). A mean realigned functional image was then produced by averaging all the realigned and unwarped functional scans. The structural T1 image was segmented into grey and white matter, using the unified segmentation approach (Ashburner & Friston, 2005) as implemented in SPM12, and then coregistered to the mean functional image. Finally, the warped functional images were spatially smoothed with an isotropic Gaussian kernel of 8 mm of full-width at half maximum (FWHM). Artifact reduction software (ART; http://www.nitrc.org/projects/artifact_detect) was used to account for motion artifact and outliers in the global mean signal intensity.

Based on Hutton et al. (2011), an in-house developed Matlab toolbox was used to extract 14 physiological regressors, obtained from a physiological noise model constructed to account for signal variability related to cardiac and respiratory phase and changes. Basis sets of sine and cosine Fourier series components extending to the 5rd harmonic for the cardiac phase and 3rd harmonic for the respiratory phase were used to model the physiological fluctuations.

For each subject, changes in BOLD responses were estimated using a general linear model in which the blocks for each MSS were modelled using boxcar functions and convolved with the canonical hemodynamic response function. Motion and physiological regressors were added in the model and considered as covariates of no interest. Correct, incorrect and no responses were separately modelled in the design matrix for each MSS and the analysis was performed only on correct responses. The design matrix also included a parametric modulator to consider reaction time. A cutoff period of 250s was used for the high-pass filter to remove low-frequency drifts from the time series, while preserving signal of interest. Due to the rapidly sampled fMRI series, the “FAST” model implemented in SPM was used to estimate serial autocorrelations (Corbin et al., 2018).

### ROI analysis

ROIs extraction was performed using the MATLAB based toolbox BRANT (BRAinNetome Toolkit) (Xu et al., 2018). We defined 3 main ROIs: (1) the dorsolateral prefrontal cortex and more particularly BA 9/46, the main region of interest (Altamura et al., 2007; Cappell et al., 2010; Haque et al., 2021; Höller-Wallscheid et al., 2017; Reuter-Lorenz et al., 2001); (2) a more extended WM network considering regions that have been implicated in age-related compensatory processes during the Sternberg task (Cappell et al., 2010; Schneider-Garces et al., 2010), including regions from the frontal lobe (BA 6, BA 9, BA 12/47, BA 44, BA 45, BA 46, BA 24, BA 32), the medial temporal lobe (the hippocampus and the parahippocampal gyrus), the parietal lobe (BA 5, BA 7, BA 39, BA 40) and the occipital lobe; (3) the thalamus; as we supposed that napping habits affect vigilance state and associated arousal regulation (Deantoni, Reyt, et al., 2024). As compensation models suggest a bilateral recruitment in both hemispheres, we distinguished left and right hemispheres in each ROI. The WM network was first divided into regions (frontal, parietal, temporal, occipital) and we then extracted individual parameter estimates of BOLD signal by memory load (2-7 letters), by ROI (DLPFC, frontal, parietal, temporal and occipital regions of the WM network and thalamus) and by hemisphere. On the whole sample, global maximum coordinates of each hemisphere by ROI were extracted by computing F-tests (p_FWE-corr_< 0.05) after application of an inclusive grey-matter mask derived from tissue probability maps (probability of voxel belonging to grey matter above 0.20) and by using the small volume correction to determine the ROI. These coordinates were then used at the individual level to determine a sphere of 30 mm in which the global maximum location was determined for each participant. From the individual’s global maximum location, a new sphere of 8 mm was created to extract the BOLD signal of significantly activated voxels (p< 0.001) by MSS. Generalised linear mixed models were conducted first to assess the effect of group (habitual nappers or non-nappers), memory load (2-7 letters), hemisphere (left or right), as well as their interactions on the extracted BOLD signal in the task-related brain region (DLPFC). The same model was computed on thalamic BOLD signal. To assess region-specificity of BOLD signal, a separate GLMM further included the factor region (frontal, parietal, temporal and occipital) and their interactions. Group, memory load, hemisphere and regions were defined as categorical fixed effects and subjects as random effects. Brain activity at each MSS was compared to the mean activity of all previous MSS (Helmert coding). Treatment contrasts were computed with non-nappers, left hemisphere and frontal lobe as references. As for behavioural data, model comparisons were performed and Akaike criterion determined that age, sex and education had not to be included as covariates. Daytime sleepiness and BMI were included as covariates as group significantly differed in these variables. Statistics were performed using R (version 3.6.3) (R Core Team, 2020).

In the context of functional compensation, hemispheric asymmetry reduction should be associated with performance maintenance (Reuter-Lorenz & Cappell, 2008). Generalised linear mixed models were thus conducted to assess whether task performance (d’ or reaction time) was predicted by WM load (2-7 letters), group (habitual nappers vs non-nappers), BOLD signal as well as their interactions. Separate models were computed for the left and right hemisphere. Group and memory load were defined as categorical fixed effects and subjects as random effects. Performance at each MSS was compared to the mean performance of all previous MSS (Helmert coding). Treatment contrast was computed with non-nappers as reference. Daytime sleepiness and BMI were added as covariates. Model comparisons were also performed and according to Akaike criterion, models assessing accuracy also include age, gender and educational level.

## Acknowledgements

The study was conducted at the GIGA-In Vivo Imaging technological platform of ULiège, Belgium. It was supported by the European Research Council (ERC, ERCStG-COGNAP) under the European Union’s Horizon 2020 research and innovation programme (Grant agreement No. 757763). This work was also supported by the Fonds de la Recherche Scientifique - FNRS under Grant nr T.0220.20. Fabienne Collette, Gilles Vandewalle, Christophe Philips and Christina Schmidt are permanent researcher at the FRS - FNRS. Michele Deantoni and Marine Dourte are FRIA grantees of the Fonds de la Recherche Scientifique - FNRS, Belgium.

## Author contributions

Christina Schmidt, Mathilde Reyt, Michele Deantoni, Vincenzo Muto, Fabienne Collette, Gilles Vandewalle and Pierre Maquet designed the study. Mathilde Reyt, Michele Deantoni, Marion Baillet, Vincenzo Muto, Alexia Lesoinne, Marine Dourte, Stella DeHaan, Sophie Laloux and Christina Schmidt collected data. Mathilde Reyt, Fabienne Collette, Mohamed Ali Bahri, Christophe Philips and Christina Schmidt developed methods, analysed and interpreted the data. All authors prepared, revised and contributed to the submitted version of the manuscript.

## REFERENCES

Akerstedt, T., & Gillberg, M. (1990). Subjective and objective sleepiness in the active individual. Int J Neurosci, 52(1–2), 29–37.

Altamura, Mario, Brita Elvevåg, Giuseppe Blasi, Alessandro Bertolino, Joseph H. Callicott, Daniel R. Weinberger, Venkata S. Mattay, and Terry E. Goldberg. (2007). “Dissociating the Effects of Sternberg Working Memory Demands in Prefrontal Cortex.” Psychiatry Research - Neuroimaging, 154(2), 103–114. 10.1016/j.pscychresns.2006.08.002.

Andersson, Jesper L.R., Chloe Hutton, John Ashburner, Robert Turner, and Karl Friston. (2001). “Modeling Geometric Deformations in EPI Time Series.” NeuroImage, 13(5): 903–919. 10.1006/nimg.2001.0746.

Ashburner, John, and Karl J. Friston. (2005). “Unified Segmentation.” NeuroImage, 26(3): 839–851. 10.1016/j.neuroimage.2005.02.018.

Bates, D., Mächler, M., Bolker, B., & Walker, S. (2015). Fitting Linear Mixed-Effects Models Using lme4. Journal of Statistical Software, 67(1). 10.18637/jss.v067.i01

Brainard, David H. (1997). “The Psychophysics Toolbox.” Spat Vis., 10(4): 433–36.

Beck, A. T., Epstein, N., Brown, G., & Steer, R. A. (1988). An inventory for measuring clinical anxiety: psychometric properties. J Consult Clin Psychol, 56(6), 893–897.

Beck, A. T., Ward, C. H., Mendelson, M., Mock, J., & Erbaugh, J. (1961). An inventory for measuring depression. Arch Gen Psychiatry, 4, 561–571.

Bennett, I. J., Rivera, H. G., & Rypma, B. (2013). Isolating age-group differences in working memory load-related neural activity: Assessing the contribution of working memory capacity using a partial-trial fMRI method. NeuroImage, 72, 20–32. 10.1016/j.neuroimage.2013.01.030.

Biesbroek, J. M., Verhagen, M. G., van der Stigchel, S., & Biessels, G. J. (2024). When the central integrator disintegrates: A review of the role of the thalamus in cognition and dementia. Alzheimer’s & Dementia, 20(3), 2209–2222. 10.1002/alz.13563.

Blackwell, Terri, Yaffe, K., Ancoli-Israel, S., Schneider, J. L., Cauley, J. A., Hillier, T. A., Fink, H. A., & Stone, K. L. (2006). Poor sleep is associated with impaired cognitive function in older women: The study of osteoporotic fractures. Journals of Gerontology - Series A Biological Sciences and Medical Sciences, 61(4), 405–410. 10.1093/gerona/61.4.405.

Buysse, D J, Reynolds, C. F., Monk, T. H., Berman, S. R., & Kupfer, D. J. (1989). The Pittsburgh Sleep Quality Index: a new instrument for psychiatric practice and research. Psychiatry Res, 28(2), 193–213. 10.165-1781(89)90047-4.

Cabeza, R, & Dennis, N. A. (2012). Frontal Lobes and Aging: Deterioration and Compensation. In D. T. Stuss & R. T. Knight (Eds.), Principles of Frontal Lobe Function (628–652). Oxford University Press.

Cabeza, Roberto. (2002). Hemispheric Asymmetry Reduction in Older Adults: The HAROLD Model, Psychology and Aging, 17(1): 85–100. 10.1037//0882-7974.17.1.85.

Cabeza, Roberto, Marilyn Albert, Sylvie Belleville, Fergus I M Craik, Audrey Duarte, Cheryl L Grady, Ulman Lindenberger, et al. (2018). Maintenance, Reserve and Compensation: The Cognitive Neuroscience of Healthy Ageing. Nature Reviews Neuroscience, 19: 701–10. 10.1038/s41583-018-0068-2.

Cappell, K A, L Gmeindl, and P A Reuter-Lorenz. (2010). Age Differences in Prefontal Recruitment during Verbal Working Memory Maintenance Depend on Memory Load. Cortex, 46(4): 462–73. 10.1016/j.cortex.2009.11.009.

Carp, J., Gmeindl, L., & Reuter-Lorenz, P. A. (2010). Age differences in the neural representation of working memory revealed by multi-voxel pattern analysis. Frontiers in Human Neuroscience, 4, 1–10. 10.3389/fnhum.2010.00217.

Carskadon, M. A., Brown, E. D., & Dement, W. C. (1982). Sleep fragmentation in the elderly: Relationship to daytime sleep tendency. Neurobiology of Aging, 3(4), 321–327. 10.1016/0197-4580(82)90020-3.

Corbin, Nadège, Nick Todd, Karl J. Friston, and Martina F. Callaghan. (2018). Accurate Modeling of Temporal Correlations in Rapidly Sampled FMRI Time Series. Human Brain Mapping, 39(10): 3884–3897. 10.1002/hbm.24218.

Davis, Simon W., Nancy A. Dennis, Sander M. Daselaar, Mathias S. Fleck, and Roberto Cabeza. (2008). “Qué PASA? The Posterior-Anterior Shift in Aging.” Cerebral Cortex, 18(5): 1201–9. 10.1093/cercor/bhm155.

Deantoni, M., Reyt, M., Baillet, M., Dourte, M., De Haan, S., Lesoinne, A., Vandewalle, G., Maquet, P., Berthomier, C., Muto, V., Hammad, G., & Schmidt, C. (2024). Napping and circadian sleep– wake regulation during healthy aging. Sleep, 47(5). 10.1093/sleep/zsad287.

Deantoni, M., Reyt, M., Berthomier, C., Muto, V., Hammad, G., De Haan, S., Dourte, M., Taillard, J., Lambot, E., Cajochen, C., Reichert, C. F., Maire, M., Baillet, M., & Schmidt, C. (2023). Association between circadian sleep regulation and cortical gyrification in young and older adults. Sleep, 46(9). 10.1093/sleep/zsad094.

Dijk, D J, and S N Archer. (2009). Circadian and Homeostatic Regulation of Human Sleep and Cognitive Performance and Its Modulation by PERIOD3. Sleep Medecine Clinics, 4(2): 111–125.

Dijk, D. J., & von Schantz, M. (2005). Timing and consolidation of human sleep, wakefulness, and performance by a symphony of oscillators. J Biol Rhythms, 20(4), 279–290.

Ding, Y., Zheng, Y., Liu, T., Chen, T., Wang, C., Sun, Q., Hua, M., & Hua, T. (2017). Changes in GABAergic markers accompany degradation of neuronal function in the primary visual cortex of senescent rats. Scientific Reports, 7(1), 14897. 10.1038/s41598-017-15006-3

Dixon, R. A., & Lachman, M. E. (2019). Risk and protective factors in cognitive aging: Advances in assessment, prevention, and promotion of alternative pathways. In The aging brain: Functional adaptation across adulthood. (217–263). American Psychological Association. 10.1037/0000143-009

Johns, M. W. (1991). A new method for measuring daytime sleepiness: the Epworth sleepiness scale. Sleep, 14(6), 540–545.

Koen, J. D., & Rugg, M. D. (2019). Neural Dedifferentiation in the Aging Brain. Trends in Cognitive Sciences, 23(7), 547–559. 10.1016/j.tics.2019.04.012

Festini, Sara B., Laura Zahodne, and Patricia A. Reuter-Lorenz. (2018). Theoretical Perspectives on Age Differences in Brain Activation: HAROLD, PASA, CRUNCH—How Do They STAC Up? Oxford Research Encyclopedia of Psychology, 1–24. 10.1093/acrefore/9780190236557.013.400.

Folstein, M. F., Folstein, S. E., & McHugh, P. R. (1975). Mini-mental state. A practical method for grading the cognitive state of patients for the clinician. Journal of Psychiatric Research, 12(3), 189–198. 10.1016/0022-3956(75)90026-6

Hammad, Grégory, Mathilde Reyt, Nikita Beliy, Marion Baillet, Michele Deantoni, Alexia Lesoinne, Vincenzo Muto, and Christina Schmidt. (2021). PyActigraphy: Open-Source Python Package for Actigraphy Data Visualization and Analysis. PLOS Computational Biology, 17(10): e1009514. 10.1371/journal.pcbi.1009514.

Haque, Zakia Z., Ranshikha Samandra, and Farshad Alizadeh Mansouri. (2021). Neural Substrate and Underlying Mechanisms of Working Memory: Insights from Brain Stimulation Studies. Journal of Neurophysiology, 126(6): 2038–2053. 10.1152/jn.00041.2021.

Höller-Wallscheid, Melanie S., Peter Thier, Jörn K. Pomper, and Axel Lindner. (2017). Bilateral Recruitment of Prefrontal Cortex in Working Memory Is Associated with Task Demand but Not with Age. Proceedings of the National Academy of Sciences of the United States of America, 114(5): E830–39. 10.1073/pnas.1601983114.

Horne, J. A., & Östberg, O. (1976). A self-assessment questionnaire to determine morningnesseveningness in human circadian rhythms. International Journal of Chronobiology, 4, 97–110.

Hua, T., Li, X., He, L., Zhou, Y., Wang, Y., & Leventhal, A. G. (2006). Functional degradation of visual cortical cells in old cats. Neurobiology of Aging, 27(1), 155–162. 10.1016/j.neurobiolaging.2004.11.012.

Hutton, C., O. Josephs, J. Stadler, E. Featherstone, A. Reid, O. Speck, J. Bernarding, and N. Weiskopf. (2011). The Impact of Physiological Noise Correction on FMRI at 7T. NeuroImage, 57(1): 101–12. 10.1016/j.neuroimage.2011.04.018.

Hutton, Chloe, Andreas Bork, Oliver Josephs, Ralf Deichmann, John Ashburner, and Robert Turner. (2002). Image Distortion Correction in FMRI: A Quantitative Evaluation. NeuroImage, 16(1): 217–40. 10.1006/nimg.2001.1054.

Kleiner, Mario, David H. Brainard, and Denis G. Pelli. (2007). What’s New in Psychtoolbox-3?. Perception, 36: 1–16.

Knights, E., Morcom, A. M., & Henson, R. N. (2021). Does hemispheric asymmetry reduction in older adults in motor cortex reflect compensation? Journal of Neuroscience, 41(45), 9361–9378. 10.1523/JNEUROSCI.1111-21.2021.

Kuznetsova, A., Brockhoff, P. B., & Christensen, R. H. B. (2017). lmerTest Package: Tests in Linear Mixed Effects Models. Journal of Statistical Software, 82(13). 10.18637/jss.v082.i13.

Leng, Yue, Susan Redline, Katie L. Stone, Sonia Ancoli-Israel, and Kristine Yaffe. (2019). Objective Napping, Cognitive Decline, and Risk of Cognitive Impairment in Older Men. Alzheimer’s & Dementia, 15(8): 1039–1047. 10.1016/j.jalz.2019.04.009.

Leng, Yue, Katie Stone, Sonia Ancoli-Israel, Kenneth Covinsky, and Kristine Yaffe. (2018). Who Take Naps? Self-Reported and Objectively Measured Napping in Very Old Women. The Journals of Gerontology: Series A, 73(3): 374–379. 10.1093/gerona/glx014.

Li, Junxin, Pamela Z. Cacchione, Nancy Hodgson, Barbara Riegel, Brendan T. Keenan, Mathew T. Scharf, Kathy C. Richards, and Nalaka S. Gooneratne. (2017). Afternoon Napping and Cognition in Chinese Older Adults: Findings from the China Health and Retirement Longitudinal Study Baseline Assessment. Journal of the American Geriatrics Society, 65(2): 373–80. 10.1111/jgs.14368.

Li, Peng, Lei Gao, Lei Yu, Xi Zheng, Ma Cherrysse Ulsa, Hui Wen Yang, Arlen Gaba, et al. (2022). Daytime Napping and Alzheimer’s Dementia: A Potential Bidirectional Relationship.” Alzheimer’s and Dementia, 19(1):158–168. 10.1002/alz.12636.

Lim, Andrew S.P., Lei Yu, Madalena D. Costa, Aron S. Buchman, David A. Bennett, Sue E. Leurgans, and Clifford B. Saper. 2011. “Quantification of the Fragmentation of Rest-Activity Patterns in Elderly Individuals Using a State Transition Analysis.” Sleep, 34 (11): 1569–1581. 10.5665/sleep.1400.

Lim, Andrew S.P., Lei Yu, Madalena D. Costa, Sue E. Leurgans, Aron S. Buchman, David A. Bennett, and Clifford B. Saper. (2012). Increased Fragmentation of Rest-Activity Patterns Is Associated with a Characteristic Pattern of Cognitive Impairment in Older Individuals. Sleep, 35(5): 633– 640. 10.5665/sleep.1820.

Lindenberger, Ulman. (2014). Human Cognitive Aging: Corriger La Fortune? Science, 346 (6209):572–78. http://science.sciencemag.org/ http://science.sciencemag.org/ http://ovidsp.ovid.com/ovidweb.cgi?T=JS&CSC=Y&NEWS=N&PAGE=fulltext&D=ovftp&AN=00007529-201410310-00032.

Luik, Annemarie I., Lisette A. Zuurbier, Albert Hofman, Eus J.W. Van Someren, M. Arfan Ikram, and Henning Tiemeier. (2015). Associations of the 24-h Activity Rhythm and Sleep with Cognition: A Population-Based Study of Middle-Aged and Elderly Persons. Sleep Medicine, 16(7): 850–855. 10.1016/j.sleep.2015.03.012.

Luik, Annemarie I., Lisette A. Zuurbier, Albert Hofman, Eus J.W. Van Someren, and Henning Tiemeier. (2013). Stability and Fragmentation of the Activity Rhythm across the Sleep-Wake Cycle: The Importance of Age, Lifestyle, and Mental Health. Chronobiology International, 30(10): 1223–1230. 10.3109/07420528.2013.813528.

Lupien, S J, and N Wan. (2004). Successful Ageing: From Cell to Self. Philos Trans R Soc Lond B Biol Sci, 359(1449):1413–26. 10.1098/rstb.2004.1516ER0XX2BW8UU7F91B [pii].

Ma, Ning, David F. Dinges, Mathias Basner, and Hengyi Rao. (2015). How Acute Total Sleep Loss Affects the Attending Brain: A Meta-Analysis of Neuroimaging Studies. Sleep, 38(2): 233–240. 10.5665/sleep.4404.

Maire, M., Reichert, C. F., & Schmidt, C. (2013). Sleep-Wake Rhythms and Cognition. Journal of Cognitive and Behavioral Psychotherapies, 13(1), 133–170.

Mattay, V. S., Fera, F., Tessitore, A., Hariri, A. R., Berman, K. F., Das, S., Meyer-Lindenberg, A., Goldberg, T. E., Callicott, J. H., & Weinberger, D. R. (2006). Neurophysiological correlates of age-related changes in working memory capacity. Neuroscience Letters, 392(1–2), 32–37. 10.1016/j.neulet.2005.09.025

Mattis, S. (1976). Mental Status Examination of Organic Mental Syndrome in the Elderly Patient. In L. Bellack & T. B. Karusu (Eds.), Geriatric Psychiaty (77–121). Grune and Stratton.

Mazerolle, M.J. (2020). AICcmodavg: Model selection and multimodel inference based on (Q)AIC(c). https://cran.r-project.org/package=AICcmodavg

McDonough, I. M., Wong, J. T., & Gallo, D. A. (2013). Age-related differences in prefrontal cortex activity during retrieval monitoring: Testing the compensation and dysfunction accounts. Cerebral Cortex, 23(5), 1049–1060. 10.1093/cercor/bhs064

Morcom, A. M., & Henson, R. N. A. (2018). Increased prefrontal activity with aging reflects nonspecific neural responses rather than compensation. Journal of Neuroscience, 38(33), 7303– 7313. 10.1523/JNEUROSCI.1701-17.2018.

Morris, Brian J., Bradley J. Willcox, and Timothy A. Donlon. (2019). Genetic and Epigenetic Regulation of Human Aging and Longevity. Biochimica et Biophysica Acta - Molecular Basis of Disease, 1865(7): 1718–44. 10.1016/j.bbadis.2018.08.039.

Muto, V., Jaspar, M., Meyer, C., Kusse, C., Chellappa, S. L., Degueldre, C., Balteau, E., Shaffii-Le Bourdiec, A., Luxen, A., Middleton, B., Archer, S. N., Phillips, C., Collette, F., Vandewalle, G., Dijk, D.-J., & Maquet, P. (2016). Local modulation of human brain responses by circadian rhythmicity and sleep debt. Science, 353(6300), 687–690. 10.1126/science.aad2993.

Nyberg, L, M Lovden, K Riklund, U Lindenberger, and L Backman. (2012). Memory Aging and Brain Maintenance. Trends Cogn Sci, 16(5):292–305. 10.1016/j.tics.2012.04.005.

Nyberg, Lars, Dahlin, E., Stigsdotter Neely, A., & Bäckman, L. (2009). Neural correlates of variable working memory load across adult age and skill: Dissociative patterns within the fronto-parietal network: Cognition and Neurosciences. Scandinavian Journal of Psychology, 50(1), 41–46. 10.1111/j.1467-9450.2008.00678.x.

Oosterman, Joukje M., Eus J.W. Van Someren, Raymond L. C. Vogels, Barbera Van Harten, and Erik J. A. Scherder. (2009). Fragmentation of the Rest-Activity Rhythm Correlates with Age-Related Cognitive Deficits. Journal of Sleep Research, 18(1): 129–35. 10.1111/j.1365-2869.2008.00704.x.

Owusu, Jocelynn T, Alexandra M.V. Wennberg, Calliope B Holingue, Marian Tzuang, Kylie D Abeson, and Adam P Spira. (2019). Napping Characteristics and Cognitive Performance in Older Adults. International Journal of Geriatric Psychiatry, 34(1): 87–96. 10.1002/gps.4991.

Papp, K. V, Rentz, D. M., Orlovsky, I., Sperling, R. A., & Mormino, E. C. (2017). Optimizing the preclinical Alzheimer’s cognitive composite with semantic processing: The PACC5. Alzheimer’s & Dementia, 3(4), 668–677. 10.1016/j.trci.2017.10.004

Park, Denise C., and Patricia Reuter-Lorenz. (2009). The Adaptive Brain: Aging and Neurocognitive Scaffolding. Annual Review of Psychology, 60, 173–196. 10.1146/annurev.psych.59.103006.093656.

Pelli, Denis G. 1997. The VideoToolbox Software for Visual Psychophysics: Transforming Numbers into Movies. Spat Vis., 10(4): 437–442.

Portas, C. M., Rees, G., Howseman, A. M., Josephs, O., Turner, R., & Frith, C. D. (1998). A Specific Role for the Thalamus in Mediating the Interaction of Attention and Arousal in Humans. The Journal of Neuroscience, 18(21), 8979–8989. 10.1523/JNEUROSCI.18-21-08979.1998.

R Core Team. 2020. “R: A Language and Environment for Statistical Computing.” Vienna, Austria: R Foundation for Statistical Computing. https://www.r-project.org/.

Reichert, C. F., Maire, M., Gabel, V., Viola, A. U., Götz, T., Scheffler, K., Klarhöfer, M., Berthomier, C., Strobel, W., Phillips, C., Salmon, E., Cajochen, C., & Schmidt, C. (2017). Cognitive brain responses during circadian wake-promotion: evidence for sleep-pressuredependent hypothalamic activations. Scientific Reports, 7(1), 5620. 10.1038/s41598-017-05695-1.

Reuter-Lorenz, P A, and D C Park. (2014). How Does It STAC up? Revisiting the Scaffolding Theory of Aging and Cognition. Neuropsychol Rev, 24(3):355–370. 10.1007/s11065-014-9270-9.

Reuter-Lorenz, Patricia A., and Katherine A. Cappell. (2008). Neurocognitive Aging and the Compensation Hypothesis. Current Directions in Psychological Science, 17(3): 177–182. 10.1111/j.1467-8721.2008.00570.x.

Reuter-Lorenz, Patricia A, Jonides, J., Smith, E. E., Hartley, A., Miller, A., Marshuetz, C., & Koeppe, R. A. (2000). Age differences in the frontal lateralization of verbal and spatial working memory revealed by PET. Journal of Cognitive Neuroscience, 12(1), 174–187. 10.1162/089892900561814.

Reyt, Mathilde, Michele Deantoni, Marion Baillet, Alexia Lesoinne, Sophie Laloux, Eric Lambot, Justine Demeuse, et al. (2022). Daytime Rest: Association with 24 - h Rest – Activity Cycles, Circadian Timing and Cognition in Older Adults. Journal of Pineal Research, e12820. 10.1111/jpi.12820.

Roe, J. M., Vidal-Piñeiro, Di., Sneve, M. H., Kompus, K., Greve, D. N., Walhovd, K. B., Fjell, A. M., & Westerhausen, R. (2020). Age-Related Differences in Functional Asymmetry during Memory Retrieval Revisited: No Evidence for Contralateral Overactivation or Compensation. Cerebral Cortex, 30(3), 1129–1147. 10.1093/cercor/bhz153.

Roenneberg, Till, Lena K. Keller, Dorothee Fischer, Joana L. Matera, Céline Vetter, and Eva C. Winnebeck. 2015. “Chapter Twelve - Human Activity and Rest In Situ.” In Methods in Enzymology, edited by Amita Sehgal, 552:257–83. Academic Press. 10.1016/bs.mie.2014.11.028.

Schiff, Nicholas D. (2008). Central Thalamic Contributions to Arousal Regulation and Neurological Disorders of Consciousness. Annals of the New York Academy of Sciences, 1129: 105–18. 10.1196/annals.1417.029.

Schmidt, C., Collette, F., Cajochen, C., & Peigneux, P. (2007). A time to think: circadian rhythms in human cognition. Cogn Neuropsychol, 24(7), 755–789.

Schmolesky, M. T., Wang, Y., Pu, M., & Leventhal, A. G. (2000). Degradation of stimulus selectivity of visual cortical cells in senescent rhesus monkeys. Nature Neuroscience, 3(4), 384–390. 10.1038/73957.

Schneider-Garces, Nils J., Brian A. Gordon, Carrie R. Brumback-Peltz, Eunsam Shin, Yukyung Lee, Bradley P. Sutton, Edward L. Maclin, Gabriele Gratton, and Monica Fabiani. (2010). Span, CRUNCH, and beyond: Working Memory Capacity and the Aging Brain. Journal of Cognitive Neuroscience, 22(4): 655–69. 10.1162/jocn.2009.21230.

Stanislaw, Harold, and Natasha Todorov. (1999). Calculation of Signal Detection Theory Measures. Behavior Research Methods, Instruments, & Computers, 31(I): 137–49.

Stern, Yaakov, Eider M. Arenaza-Urquijo, David Bartrés-Faz, Sylvie Belleville, Marc Cantilon, Gael Chetelat, Michael Ewers, et al. (2020). Whitepaper: Defining and Investigating Cognitive Reserve, Brain Reserve, and Brain Maintenance. Alzheimer’s and Dementia, 16(9): 1305–11. 10.1016/j.jalz.2018.07.219.

Sternberg, S. (1966). High-speed scanning in human memory. Science, 153(3736), 652–654. 10.1126/science.153.3736.652.

Sturm, W., A. De Simone, B. J. Krause, K. Specht, V. Hesselmann, I. Radermacher, H. Herzog, L. Tellmann, H. W. Müller-Gärtner, and K. Willmes. (1999). Functional Anatomy of Intrinsic Alertness: Evidence for a Fronto-Parietal-Thalamic-Brainstem Network in the Right Hemisphere. Neuropsychologia, 37(7): 797–805. 10.1016/S0028-3932(98)00141-9.

Sun, J., Ma, C., Zhao, M., Magnussen, C. G., & Xi, B. (2022). Daytime napping and cardiovascular risk factors, cardiovascular disease, and mortality: A systematic review. Sleep Med Rev, 65. 10.1016/j.smrv.2022.101682

Taillard, J., Gronfier, C., Bioulac, S., Philip, P., & Sagaspe, P. (2021). Sleep in Normal Aging, Homeostatic and Circadian Regulation and Vulnerability to Sleep Deprivation. Brain Sciences, 11(8). 10.3390/brainsci11081003.

Wickham, H., Averick, M., Bryan, J., Chang, W., McGowan, L., François, R., Grolemund, G., Hayes, A., Henry, L., Hester, J., Kuhn, M., Pedersen, T., Miller, E., Bache, S., Müller, K., Ooms, J., Robinson, D., Seidel, D., Spinu, V., … Yutani, H. (2019). Welcome to the Tidyverse. Journal of Open Source Software, 4(43), 1686. 10.21105/joss.01686.

Weissová, Kamila, Aleš Bartoš, Martin Sládek, Marta Nováková, and Alena Sumová. (2016). Moderate Changes in the Circadian System of Alzheimer’s Disease Patients Detected in Their Home Environment. PLoS ONE, 11(1): 1–19. 10.1371/journal.pone.0146200.

Xu, Kaibin, Yong Liu, Yafeng Zhan, Jiaji Ren, and Tianzi Jiang. (2018). BRANT: A Versatile and Extendable Resting-State FMRI Toolkit. Frontiers in Neuroinformatics, 12: 1–13. 10.3389/fninf.2018.00052.

Zuurbier, Lisette A., Annemarie I. Luik, Albert Hofman, Oscar H. Franco, Eus J.W. Van Someren, and Henning Tiemeier. (2015). Fragmentation and Stability of Circadian Activity Rhythms Predict Mortality. American Journal of Epidemiology, 181(1): 54–63. 10.1093/aje/kwu245.

